# Tunable fluorogenic DNA probes drive fast and high-resolution single-molecule fluorescence imaging

**DOI:** 10.1101/2025.01.21.634148

**Authors:** Mirjam Kümmerlin, Qing Zhao, Jagadish Hazra, Christof Hepp, Alison Farrar, Piers Turner, Achillefs N. Kapanidis

## Abstract

A main limitation of single-molecule fluorescence (SMF) measurements is the “high concentration barrier”, describing the maximum concentration of fluorescent species tolerable for sufficient signal-to-noise ratio (SNR). To address this barrier in several SMF applications, we design fluorogenic probes based on short ssDNAs, fluorescing only upon hybridising to their complementary target sequence. We engineer the quenching efficiency and fluorescence enhancement upon duplex formation through screening several fluorophore-quencher combinations, label lengths, and sequence motifs, which we utilise as tuning screws to adapt our labels to different experimental designs. Using these fluorogenic probes, we can perform SMF experiments at concentrations of 10 µM fluorescent labels; this concentration is 100-fold higher than the operational limit for standard TIRF experiments. We demonstrate the ease of implementing these probes into existing protocols by performing super-resolution imaging with DNA-PAINT, employing a fluorogenic 6 nt-long imager; through the faster acquisition of binding events, the imaging of viral genome segments could be sped up significantly to achieve extraction of 20-nm structural features with only ∼150 s of imaging. The exceptional tunability of our probe design will overcome concentration barriers in SMF experiments and unlock new possibilities in super-resolution imaging, molecular tracking, and smFRET.

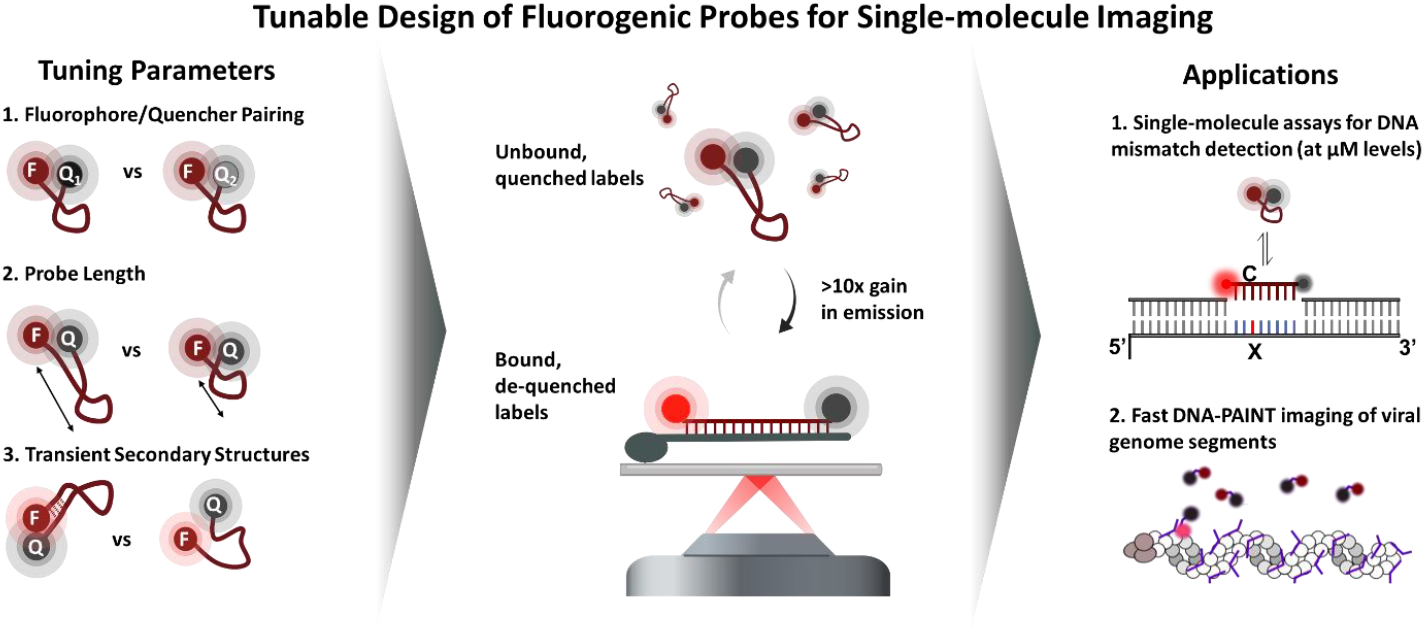

## Introduction

The study of structure, dynamics, and function of biological systems has been transformed by the unique insights gathered from their study using single-molecule methods both *in vitro* and *in vivo*.^1^ Their versatility and sensitivity have made single-molecule fluorescence (SMF) spectroscopy and microscopy studies especially popular, particularly when compatibility with use in living cells is required.^2,3^The detection of a fluorescing molecule, its emission properties, and its precise location enable measurements of molecular stoichiometries, as well as super-resolution imaging and single-molecule tracking.^4–10^

In many SMF experiments, a critical limitation known as the “high concentration barrier” complicates single-molecule analysis when the concentration of fluorescent species exceeds a certain concentration (1 – 10 nM for general wide-filed microscopy, 100 nM for total internal reflection (TIRF) microscopy, and 50-100 pM for solution-based confocal microscopy).^11–13^ At higher concentrations of fluorescent molecules, the signal-to-noise ratio (SNR) deteriorates too far to allow for a reliable analysis of the data. This limitation introduces the need to remove any unbound or non-specifically bound fluorophores (e.g., in immunohistochemistry- or Halo-Tag-based labelling approaches ^14,15^) and to limit experimental designs with correctly labelled species to relatively low concentrations, which are often far below biologically relevant affinities.^12,13^

Strategies to address the concentration barrier have focused both on limiting the excitation of or the emission from unbound (or otherwise diffusing) fluorescent molecules in the background, and on enhancing the fluorescence of molecules involved in the process or bound to the structures of interest. Background suppression can be achieved by suitably modified instruments, such as in zero-mode-waveguides (ZMWs), using the evanescent excitation field in a TIRF microscope or via the employment of a second laser inducing stimulated-emission depletion (STED) in STED-microscopy, which reduces the de-facto excited volume. The standard for SMF experiments, the TIRF microscope, however, still cannot allow routine detection of single immobilised molecules in the presence of >100 nM of fluorescent species.^12,16–18^

Fluorescent enhancement of only those molecules which are under observation using plasmonic effects can improve the SNR of a measurement by several orders of magnitude^19^, but require the experimental protocol to be compatible with the stability, accessibility and functionality offered by the specific nanostructure employed in the experiment. As such, nanostructures facilitating plasmonic enhancement are so far unsuitable for imaging or tracking experiments.

A third approach focuses on using fluorogenic molecules which exhibit a significant increase in quantum yield/fluorescence emission upon interacting with their target structure. Examples of this include fluorogenic dyes (e.g., Nile red in PAINT microscopy and different intercalators ^20–23^) or molecular structures in which a conformational change upon binding displaces a dark quencher (Q) to recover fluorescence of a fluorophore (F).

In particular, DNAs have a long-standing history of being used as fluorogenic sensors. A main focus of the early development has been on PCR probes, allowing detection and quantification of PCR products without further post-processing.^24,25^ In early days, the read-out relied on either nuclease digestion of the ssDNA to remove a Q from the proximity of a F or free the F into solution.^24–26^ Fluorogenicity upon hybridisation of these ssDNAs was first used in molecular beacons, which use a self-complementary stem region to facilitate interactions between terminally attached F and Q.^27,28^ Later on, this secondary structure was found to not be required for fluorogenicity of ssDNAs.^29^ The utilised quencher molecules have been diverse: fluorescent dyes, dark quenchers, DNA bases (with their intrinsic quenching ability), and metal nanoparticles have been employed. ^24,25,27,29–31^

The quenching process in such short doubly-labelled ssDNAs can contain FRET-quenching and contact-quenching components, both of which show different spectral and molecular dependencies.^29,32,33^ Many studies have focused on generating high quenching efficiencies, with little focus on the brightness of the de-quenched construct after hybridisation. Yet, when translating the concept from ensemble into single-molecule settings, the de-quenched state is particularly important, since a single emitter must be bright enough to be detected or localised. The deployment of fluorogenic probes in SMF and localisation-based SRM has thus been limited, requiring significant protocol and sequence engineering and limiting the probe length to above 15 nt.^34–37^

In contrast to other approaches that address the concentration barrier, the employment of these fluorogenic labels or components does not require any specialised optics or nanofabrication - but is compatible with either.^38^

Here, we present a general strategy how to design fluorogenic ssDNA-based probes based on terminal labelling with F and Q, which can be employed in a wide range of experimental set-ups, biological processes, and biomolecules. Through careful characterization of the effect the main parameters in the design of such probes (F-Q pairing, probe length, and sequence effects), we address previous limitations for short probes (5 - 15 nt). The strategy we devise allows for “plug and play” addition of fluorogenicity into experimental protocols, which we demonstrate in DNA-PAINT imaging using a 6-nt fluorogenic imager. Furthermore, we can demonstrate operational SMF assays at micromolar concentrations, highlighting the wide range of possible experimental implementations of our findings.

## Methods

### Characterisation experiments

#### Probe design

All labelled probe sequences can be found in Table S1. DNA Probes were modified to carry a 5’-ATTO647N and a 3’-Q. Probes of different lengths were designed to conserve the 3 nt surrounding the F and Q attachment site (5’-ATTO647N-TTT-X_n_-TTT-Q), with only the central section altered to facilitate different lengths. The 18 nt probe which contained a 4 nt self-complementary region differs from this design. All probes were ordered from biomers.net (Ulm, Germany). Complementary, biotinylated docking strands were ordered from Merck.

#### Sample preparation

Target strands were immobilised via a biotin binding NeutrAvidin on coverslips coated by polyethylene-glycol (PEG). In wells of silicone gaskets, 20 µl of the target strands (100-500 pM) were incubated for 10-30 s, followed by washing three times with 200 µl PBS. Subsequently, 30 µl of DNA imaging buffer (200 mM MgCl_2_, 10 mM NaCl, 50 mM HEPES pH 7.4, 6 mM BSA, 3 mM TROLOX, 1 % Glucose, 40 µg/ml catalase and 0.1 mg/ml glucose oxidase) containing the stated probe concentrations were added.

#### Imaging

SMF movies were collected using the Nanoimager SMF microscope (Oxford Nanoimaging) in objective-based TIRF illumination mode, with the excitation angle set at 54.0°. We performed the imaging using continuous-wave excitation (640 nm, with the laser powers of 9% (1.4 mW)). In all experiments, we used 100 ms exposures.

#### Image processing

To obtain brightness information for localised spots (‘signal’), all ‘.tif’ filed obtained from imaging at 10nM were processed in the Picasso ‘localize’ tool^39^. Photon counts of these localisations were frequency-counted and distributions were fitted to obtain mean photon counts for each probe. In some samples, we observed two brightness populations. We excluded the population resulting from probes without functional quenchers (either bleached Qs or faulty synthesis), which exhibits the same brightness as probes with only the F labelling. The obtained photon counts (S_FQ_) were normalised by dividing them by the average photon counts observed for probes without Qs (S_F_) to obtain the ‘normalised signal’ S_N_.

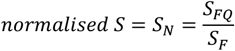

From all videos, background intensities were extracted using a custom script, averaged over the central area of the field of view and across the first 10 frames of the movies. These background intensities were plotted against the probe concentration and fitted with a linear regression in Origin (Origin Labs) to extract the molar background B_FQ, mol_. These were normalised by dividing them by the molar background emission of probes only labelled with a F (B_F,mol_) to calculate the ‘normalised background’, B_N,mol_.

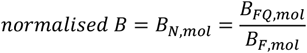

The normalization applied here ensures that our evaluation is not sensitive to e.g., changes in quantum yield of the dye when going from ss to ds state, and instead only captures the changes induced by the quencher being present. The Fluorogenic Factor (FF) capturing the overall fluorogenicity of a probe, was calculated for all samples by dividing the normalised signal by the normalised background:

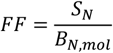

### Single-molecule hybridisation assay

#### Sample preparation

Oligos were obtained from biomers.net and Merck, dissolved to a final concentration of 100 μM, and stored at -20ºC (for Sequences, see Table S1). All components of the gapped target were mixed in annealing buffer (200 mM Tris–HCl pH 8.0, 500 mM NaCl, 1 mM EDTA) at 2 μM, and then annealed in a thermocycler (program: heating to 90°C, then cooling to 25°C at 2°C/min, storing at 4°C). The labelled 8 nt probe was ordered from biomers.net (Ulm, Germany), carrying a 3’ Atto647N and a 5’-BHQ1. Preparation of surfaces and immobilisation was performed as described for characterisation experiments. Imaging was performed in buffer containing 20 mM HEPES, 200 mM NaCl, 80 mM MgCl_2_, 5% dextran sulphate, and 5% formamide.

#### Imaging

Samples were imaged at 2 μM under the same conditions described for the characterisation experiments. Imaging was performed at 1.4 mW of 640 nm and 0.8 mW of 532 nm excitation with an exposure time of 200 ms.

#### Data analysis

Molecules were localised in the green channel, traces were extracted and fitted with an HMM using custom Python software. Binding kinetics were fitted using the probability density functions correcting for finite exposure times of the camera and length of states in MEMLET.^40^ All graphs were plotted in Origin (Origin Labs).

### Viral DNA-PAINT imaging

#### Virus Strain

A/Puerto Rico/8/34 (H1N1) influenza virus (PR8) purified from embryonated chicken eggs was purchased from Charles River Laboratories, aliquoted, flash-frozen in liquid N_2_ and stored at\ -80 °C before use.

#### Probe design

Primary probes against the NA and the PB1 segment were designed as outlined in Hepp *et al*.,^*41*^ the PB1 probes were adopted to contain multiplexed docking sites against the 6-nt imager (5’-CCACCACCACCA-3’). The P3 imager sequence was used as described previously^39^ and was fluorescently labelled with Cy3B on the 5’ end. The 6nt imager has the following sequence: 5’-TGGTGG-3’ and was labelled with ATTO643 on the 5’ and with BMNQ1 on the 3’ end. Imager strands were purchased from metabion international AG (P3-Cy3B, Planegg, Germany) or biomers.net (6nt imager, Ulm, Germany).

#### Virus sample preparation

All virus samples were prepared as outlined in Hepp *et al*.^*41*^. Cover slips were cleaned (sonic bath and plasma cleaner) to remove any contaminations immediately before use. Viral particles were thawed on ice and fixed using 4 % formaldehyde. After dilution in 0.9 % NaCl solution, viral particles were added into CultureWell gaskets (Ø 6 mm, Grace Biolabs, US) mounted on the cleaned coverslips, and the liquid evaporated at 34°C. After removal of the gaskets, sticky slides VI0.4 (ibidi, Germany) were used to create flow channels over the viral particles. Viral particles were then permeabilised with 0.5 % Triton-X-100 before primary probes were added to a final concentration of 4 µM and incubated over night at 37°C. Unbound probes were removed using a clearing buffer before DNA-PAINT imaging was performed in imaging buffer (buffer B+^39^ with 75mM MgCl_2_, supplemented with 3 mM Trolox, 1 % Glucose, 40 µg/mL catalase and 0.1 mg/mL glucose oxidase).

#### Imaging

All imaging steps were performed as described in the characterisation section at laser powers of 3.6 mW (532 nm) and 150 mW (635 nm) at exposure times of 200 and 20 ms, respectively.

#### Image Processing

The image processing steps are following those outlined in Hepp *et al*.,^41^ adjusted to adopt the different noise behaviour in these experiments. Localizations were detected with Picasso “Localize”^39^ with the following sections: box length: 7, gradient: 3000, baseline: 400, sensitivity: 2.75, quantum efficiency: 0.82, pixel size: 117 (nm). The gain (= sensitivity * quantum efficiency) and offset (= baseline) are estimated using the sCMOS analysis provided by GDSC imageJ plugin.^42^ Localisations were then undrifted using the AIM algorithm^43^ implemented in picasso render using the following parameters: Segmentation: 100 frames, Intersection distance: 20 nm, max. drift in segment: 60 nm. DBSCAN^44^ was performed to find clusters with the following parameters: eps: 0.5 pixel, min_samples: 0.01 * frame number, rounded to integers. Clusters were then filtered by core sample ratio (number of core samples / total samples > 0.8) and with component number = 2.

To extract physical features from the found clusters, each localization in a cluster is rendered as a Gaussian with a sigma of 3.9 nm, then a low pass filter (order-one Butterworth filter with a cut-off frequency of 0.02) and Gaussian filter (sigma 5.85 nm) are applied to the image. Next, each localisation is transferred to a binary image using the ISODATA method^45^ excluding any clusters larger than 5000 nm^2^, then the Hilditch skeleton^46^ is found to represent the segment (“spine”). If branches exist in the skeleton, the longest one is chosen. The median value of the distance between each pixel on the skeleton and the boundary of the binary image is considered the radius of the segment. Reconstructed super-resolution images were exported from Picasso “render”.^39^

## Results

### Tuning the properties of self-quenching ssDNA probes using fluorophores and dark quenchers

To generate fluorogenic probes, we focused on doubly labelled ssDNAs, which are well suited for labelling in a lab environment for a variety of reasons. First, the sequence allows for specific interactions even at very short lengths (see DNA-PAINT applications using <10 nt long ssDNAs^39^) and provides versatility for multiplexing.^36,47,48^ Further, the strength of interaction (and consequently the kinetics of hybridization) can be tuned across a wide range of time scales (from very transient interactions in DNA-PAINT microscopy (100s of ms^48^) to fluorescent labelling of DNA origami structures which can be stable for weeks^49,50^). ssDNA oligos are widely available from a range of commercial suppliers, making the technology available to and affordable for researchers outside of expert synthesis laboratories. The spectral properties of the probe can be varied by the choice of the conjugated fluorophore, of which a huge variety is commercially available. DNAs are compatible with live-cell experiments (especially once protected by chemical modifications), and do not change the size of the targeted molecule significantly, leaving properties such as molecule diffusion of macromolecular target unaffected.

The generation of fluorogenicity in our probes follows a simple principle: a ssDNA is labelled terminally with a fluorescent dye on the 5’ end, and a dark quencher on the 3’ end. When in solution, the short persistence length of ssDNA (≈ 1 nm^51^) allows it to collapse into a random coil, whereby dye and quencher are kept in close enough proximity to allow for contact-mediated or high-efficiency FRET quenching of the dye’s fluorescence (Figure 1A). Upon hybridisation, the dsDNA duplex, due to its much higher persistence length (≈ 50 nm), separates dye and quencher, de-quenching the probe and increasing fluorescence emission. This concept has been adopted in many applications,^34,35,37,52^ but it was so far assumed that a useful contrast between dark and bright state would require the DNA to be chosen from a limited range of lengths (15-20 nt) with little possibility to apply the same fluorogenic strategy to shorter or longer constructs without loss of performance. The underlying assumptions were that, in a shorter probe, the remaining FRET quenching would not allow for bright enough fluorescence in the bound state, while for longer constructs the quenching in the unbound state would not be efficient enough to allow for significant fluorogenicity, unless hairpin structures were designed into the probes, as for example in molecular beacons.^29,36,53^

**Figure 1:**
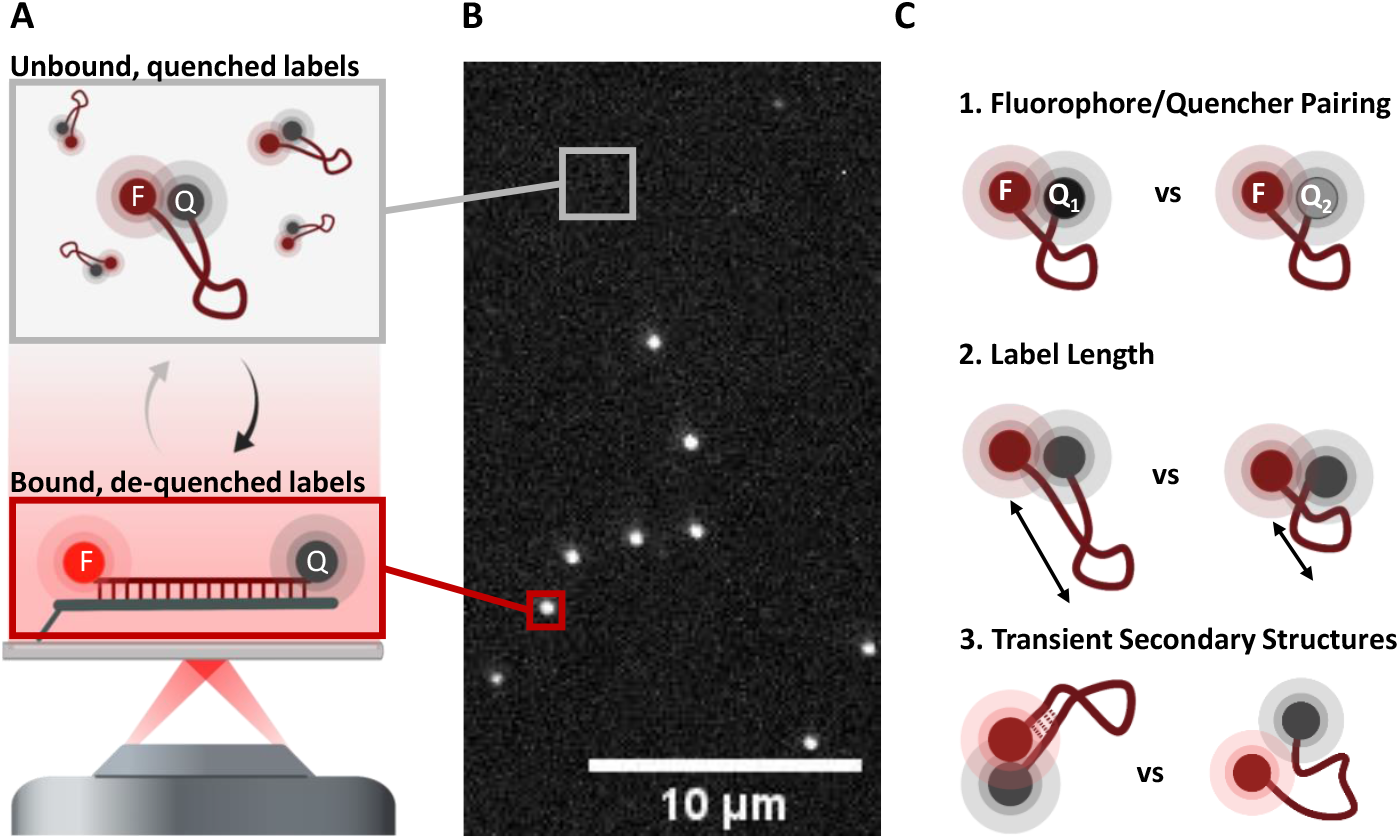
Experimental set-up for characterization experiments: **A:** ssDNA probes carry a fluorophore (F) and dark quencher (Q) on the 5’ and 3’ ends, respectively. In the single-stranded state in solution, the fluorescence emission is quenched. Upon binding their complementary docking strand immobilized on a surface, the fluorescence emission increases due to the increased distance between F and Q. **B:** The measured intensity of spots (‘signal’, red square) and the background (grey square) is extracted from the raw video and compared to values of probes only carrying the F. **C:** We evaluate three main parameters of the probes, the R_0_ of the F/Q pairing, the length of the label, and the effect of transient secondary structures.

Here we show that through controlling and tuning the emission of the probes in the quenched and de-quenched state, we can generate fluorogenic probes that overcome the previous design limitations and push boundaries in a wide range of applications, including super-resolution imaging, and SMF assays.

The basis for the fluorogenicity of our labels is the FRET interaction between F and Q, and potential further contributions by contact quenching in the single-stranded state.

In the following, we characterise three key design features our labels (Figure 1C): First, the **R**_**0**_ of the F/Q pair is reflecting the strength and spatial sensitivity of the FRET interaction. Second, the **length** of the ssDNA will determine the distance changes between F and Q upon hybridisation. Third, the **DNA sequence** determines the specificity of target binding and influences the hybridisation kinetics.

Further, internal stabilisation through **transient secondary structures** can be an important factor in quenching background fluorescence of unbound probes. We, however, don’t want to rely on the these interactions for high fluorogenicity: Some experimental designs might demand the a specific sequence while others will more crucially rely on fast kinetics of probe binding (e.g., DNA-PAINT imaging). Then it is crucial to eliminate secondary structures and internal interactions between bases e.g., by reducing the number of bases to two non-complementary ones.

To systematically assess the influence of these parameters (R_0_, length, structural elements), we look at the performance of a set of test or reference probes. These probes were designed to conserve the immediate environment of the dye and to be free of any secondary structure (by only consisting of two non-complementary bases).

The effects of these parameters were characterised by calculating the fluorogenic factor (FF) for each probe, which compares the emission in the bound (‘signal’) and unbound state (‘background’) of the labels in the typical setup of single-molecule TIRF experiments (Figure 1A-B, for more details, see methods).

Both signal and background are normalised to the levels observed for F-only probes, this normalisation removes any dependence on fluorescence changes observed upon binding of F-only probes (e.g., due to changes in quantum yield of a F when forming dsDNA) and extracts the effect the addition of a Q is having. We calculate the fluorogenic factor FF, which is the fold change in emission intensity upon hybridisation:

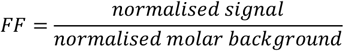

### Influence of the Förster radius of the fluorophore/quencher pair on fluorogenicity

In the quenched state, the probe design is relying on contact-quenching or very high-efficiency FRET-based quenching of the dye fluorescence. Both require close proximity and at least a minimal spectral overlap.^54^ Upon target binding, the increased distance between F and Q allows for dequenching and recovery of the fluorescence signal. Any remaining quenching in the bound state is due to FRET for distances which exceed the range of any contact quenching (assumed to be the case for all tested probes here).

For the FRET component, the spectral overlap between F emission and Q absorption is one of the factors in determining the Förster radius R_0_. Here, we thus tested several different Qs on the 3’ end of a 15-nt long construct labelled with a 5’ F (Figure 2). Based on spectral measurements and extinction coefficients, we estimated their Förster Radii to be 3.5 nm, 5.2 nm, and 6.2 nm for BHQ1, BHQ2, and BBQ650, respectively, when paired with Atto647N as F. We chose 15 nt as probe length to allow for reliable detection of both signal and background fluorescence (both well above instrument noise levels in all cases). To minimise any effects of structure-based changes in quenching behaviour, all quenchers were chosen to have similar molecular structures.

**Figure 2:**
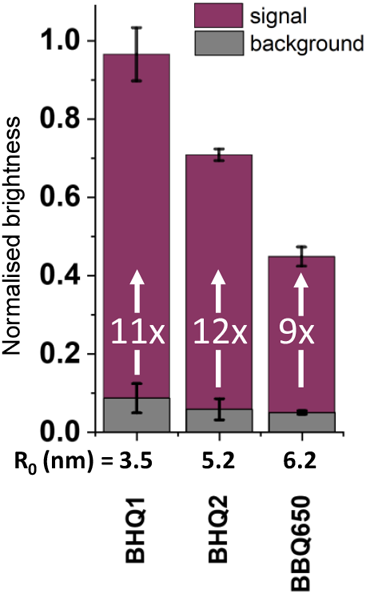
The effect of varying spectral properties of the quencher on fluorogenicity. **Red:** Signal intensity in the hybridised state, normalised to the F-only reference construct. With increasing R_0_ of the F/Q pair, the recovered signal from hybridised probes decreases. **Grey:** Background emission by unbound probes, normalised to the F-only reference construct. The background emission is efficiently quenched in all tested constructs, and higher for F/Q pairs with large R_0_. The FF as the measure for overall performance is similar for all constructs, with the BBQ650 performing worse (FF = 8.8 ± 0.2) than both BHQs (FF = 11.9 ± 0.4 and FF = 11.1 ± 0.9). Mean values and standard deviation from three independent experiments.

In the quenched, single-stranded state, the background is decreasing with increasing R_0_ of the F/Q pair, consistent with a higher degree of FRET quenching (Figure 2, ‘background’ in grey).

However, even in the case of the lowest R_0_ (3.5 nm, ATTO647N and BHQ1), we find efficient quenching, with a background of below 10 % of the F-only reference construct.

The fluorescence signal upon probe hybridisation follows the same trend (Figure 2, ‘signal’ in red): a large R_0_ correlates with low signal from the de-quenched probe. With the ATTO647N-BHQ1 construct, the full signal can be recovered, whereas for the highest spectral overlap construct (with BBQ650), we recover ≈ 45 % of the signal of the dye-only reference. For BHQ2 with the intermediate R_0_, the recovered signal is ≈ 70%.

Our results highlight the quencher choice has to be carefully balanced for an optimal fluorogenic probe. In the unbound state, a quencher with large spectral overlap (e.g., BBQ650 for ATTO647N) is most desirable; in contrast, for the bound and de-quenched state, probes with a small spectral overlap between dye and quencher have the brightest signal.

Upon calculation of the FF, we can see that the constructs with BHQ2 and BHQ1 perform best (FF = 11.9 ± 0.4 and FF = 11.1 ± 0.9, respectively), with BBQ650 achieving a slightly lower FF of 8.8 ± 0.2. The small differences suggest that high FFs can be achieved with a range of quenchers and that the choice of quencher allows to dim or brighten a specific probe in both states. We anticipate that the FF of a particular label depends significantly on how the distance changes align with steep parts of the distance-vs-FRET-Efficiency curve, and might thus be more significant in probes of different length (shorter or longer than 15nt).

Ultimately, the R_0_ thus serves as a “tuning screw” in the experimental design: an assay which requires a very short probe (e.g., for fast DNA-PAINT imaging), i.e. where even in the de-quenched state F and Q are very close, a low spectral overlap can help to recover enough signal to perform insightful measurements. On the other end of the length spectrum, probes where hybridisation allows for a large separation of F and Q, the background can be more efficiently suppressed than with a F/Q-pair with a high R_0_. The ideal quencher thus always depends on the specific requirements of the experiment and the choice should be made taking those into account.

### Influence of the probe length on the flurogenicity

In a distance-sensitive process such as FRET, the probe length has a profound impact on the fluorogenicity. We characterised a series of probe lengths spanning across the range of 5 to 25 nt, with 5 nt being the shortest we were able to observe hybridisation to surface-immobilised complementary DNA. We chose a F/Q pair with low spectral overlap (ATTO647N and BHQ1) to allow for sufficient de-quenching of the very short probes to be able to reliably detect them. Figure 3 shows the observed signal and background emission (Figure 3B) for probes of different lengths, from which we calculated the FF (Figure 3A).

**Figure 3:**
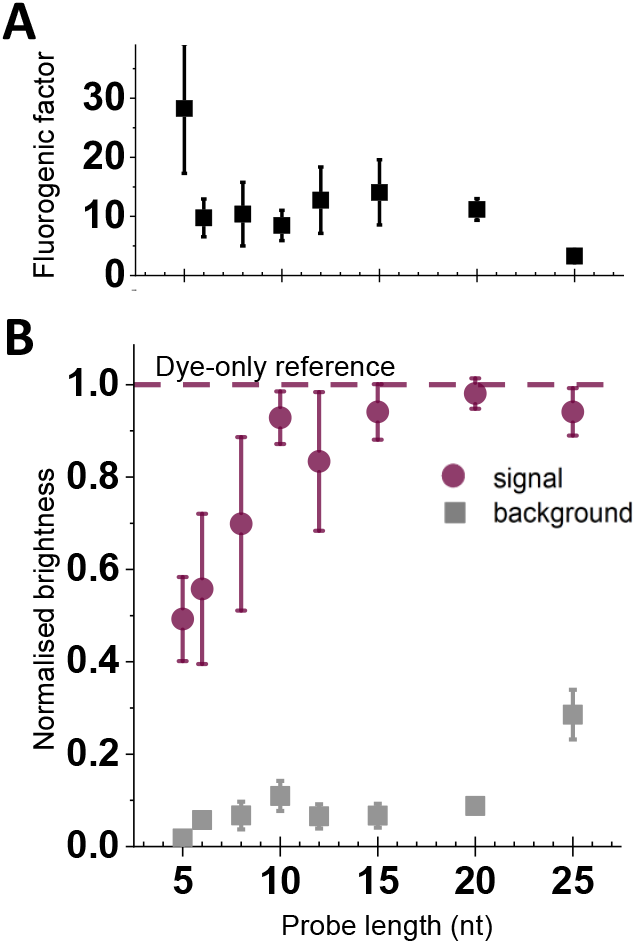
Length dependency of Fluorogenicity. **A:** The fluorogenic factor (FF) calculated as the ratio of signal and background in panel B. The 5 nt probe showed the highest fluorescence enhancement upon hybridisation, between 6 and 20 nt, the FF was constant, it then decreases for longer probes. **B: Red:** Signal intensity in the hybridised state, normalised to the dye-only reference construct. With increasing length, the signal increases and saturates from 10 nt onwards at the intensity which is expected for a case without any residual FRET-quenching. **Grey:** Background emission by unbound probes, normalised to the dye-only reference construct. The background emission is efficiently quenched in all tested constructs with the lowest background level observed with the 5 nt probe. Background in general increases with label length, with a stable level for intermediate lengths between 8 and 20 nt. Mean values and standard deviation from three independent experiments.

Overall, the background increases with increasing length, from the 5 nt probe (which emits only ≈1% of the F-only probe) to the 25 nt probe (which emits ≈27% of the unquenched probe; Figure 3B). However, for lengths between 6 and 20 nt, the background shows no significant increase, suggesting that the dynamics of the structure allow for a similar level of quenching across these lengths. Even the 25-nt probe is still 73% quenched, presumably due to its collapse into a random coil, which allows the dye and quencher to reach distances on average well below the R_0_ of the F/Q pair (3.5 nm, see Figure S1).

In the de-quenched state, the signal intensity follows the expectation for FRET-based quenching at the distances determined by the probe length. Most promisingly, the short probes (5-8 nt) show an emission of 50-70% of the saturation fluorescence signal (Figure 3B) and can clearly be imaged in our SMF set-up. Using a FRET pair with a small R_0_, as in this case, has allowed us to recover sufficient fluorescence emission in the de-quenched state for single-molecule measurements, despite the very short distance in the dsDNA state formed upon hybridisation. With increasing length, the signal further increases, until it saturates at the full fluorescence level from 10-12 nt onwards (where distances approach and then exceed 2xR_0_).

Upon calculating the FF (Figure 3A), we see that the shortest probe (5 nt) has the highest FF, mainly caused by very low background levels, potentially hint at additional quenching processes beyond the ones seen in longer probes (or the average distance of dye and quencher is much lower than in longer probes), resulting in very efficient FRET-quenching.

There is a length window between 10 and 20 nt, where (almost) the full signal is recovered and there is still efficient background suppression, leading to a FF of ≈10 for all lengths. For the 25 nt label, the increased background reduces the FF and we would thus expect to see even lower level of fluorogenicity from this F/Q pair for probes exceeding lengths of 25 nt.

The described observations were made for this specific F/Q pair. For a more efficiently quenching pair (with a larger R_0_), we expect the signal in the hybridised state to only be fully recovered at longer lengths, i.e., greater distances between F and Q. For the ssDNA state, the background increase might then also only occur for longer probes, making this a promising approach for the design of longer fluorogenic probes. On the contrary, for very short probes, the R_0_ should be chosen to be as short as possible to allow for sufficient signal recovery upon hybridisation.

### Transient secondary interactions to further reduce background fluorescence

Based on the strong distance-dependency of the FF, we hypothesised the introduction of transient secondary interactions could help further improve the FF, especially in longer probes (Figure 4A). This approach is used in molecular beacons, which often use hairpin-like structures created by the complementarity within a stem region, usually situated at either end of the probe. Often, these stem regions fairly long (e.g., 6 base-pairs^28^), forming relatively stable structures at most experimental conditions. Consequently, the hybridisation kinetics may be affected significantly, which can render the probe unsuitable for certain applications.

**Figure 4:**
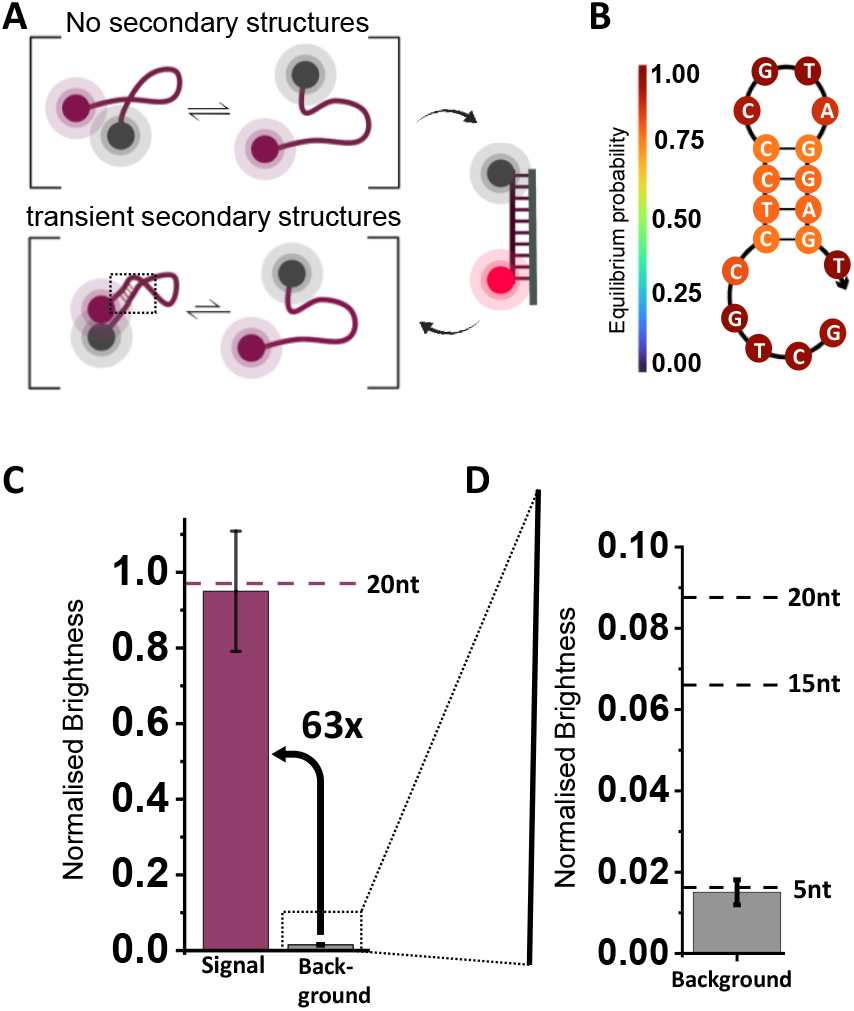
Secondary structures can improve fluorogenicity. **A:** Schematic of the reduced dynamics in the unbound, ssDNA state of probes capable of forming intramolecular basepairs, and leading to an reduced average F-Q distance. **B:** Nupack prediction of secondary structure at 25°C based on a 4-nt self-complementary region of the 3’- and center section of the sequence. **C:** Signal intensity in the hybridised state and background level, normalised to the dye-only reference construct. The 18-nt probe with secondary structure emits the same signal level as the F-only reference and 20-nt sample, suggesting full hybridisation and no residual FRET quenching. The background is reduced to 1.5 - 2%, resulting in an FF = 63, 5 x larger than in reference constructs of comparable lengths. **D:** Zoom-in of panel C. Background emission by unbound probes, normalised to the dye-only reference construct. Secondary structure in the 18-nt probe is significantly reducing the background emission compared to probes of similar length without any secondary structures (as indicated by reference lines). Mean values and standard deviation from three independent experiments.

Here, we focus on a short 4-nt intramolecular complementarity within an 18nt probe (×Delta;G = -4.23 kcal/mol at 25°C based on NUPACK^55^). The NUPACK-prediction shows that this is not forming a terminal stem region, but rather folds the 3’ end into the centre region (Figure 4B). We hypothesized that this would shift the equilibrium of possible ssDNA states in the unbound state towards those with base-pairing, and hence impose a closer F-Q distance (Figure 4A). At the same time, the stability of this interaction is low, and the remaining 5’ segment of the probe can serve as a toehold (REF) to facilitate efficient binding to a target.

In our single-molecule experiments, the signal upon hybridisation was not affected by this modification, since we recovered ≈ 100% of the fluorescence signal (Figure 3C). At the same time, we could indeed confirm that even a 4-nt complementarity was sufficient to reduce the background emission of this 18-nt probe to a similar level observed in the (much shorter) 5-nt probe (Figure 3D). Consequently, the FF of ≈ 63 in this probe was the highest we have observed in any of the constructs, and much higher than comparable probes without secondary structures exhibit (compared with 15- and 20-nt probes in Figure 3A).

This approach should be extendable to any long probe, where a complementary region of a certain length can bring dye and quencher into close proximity, even without a stable structure. Overhanging single-stranded regions can help with hybridisation kinetics.

### High-concentration single-molecule experiments

The design parameters we have characterised allow for the generation of highly fluorogenic probes which can be used in single-molecule experiments at concentrations orders of magnitude above the concentration limit of 100 nM. For example, use of the 18-nt probe with a transient secondary structure (Figure 4) allows clear single-molecule detection at concentrations up to 10 µM (Figure 5A) along with full downstream analysis for brightness, location and other properties.

**Figure 5:**
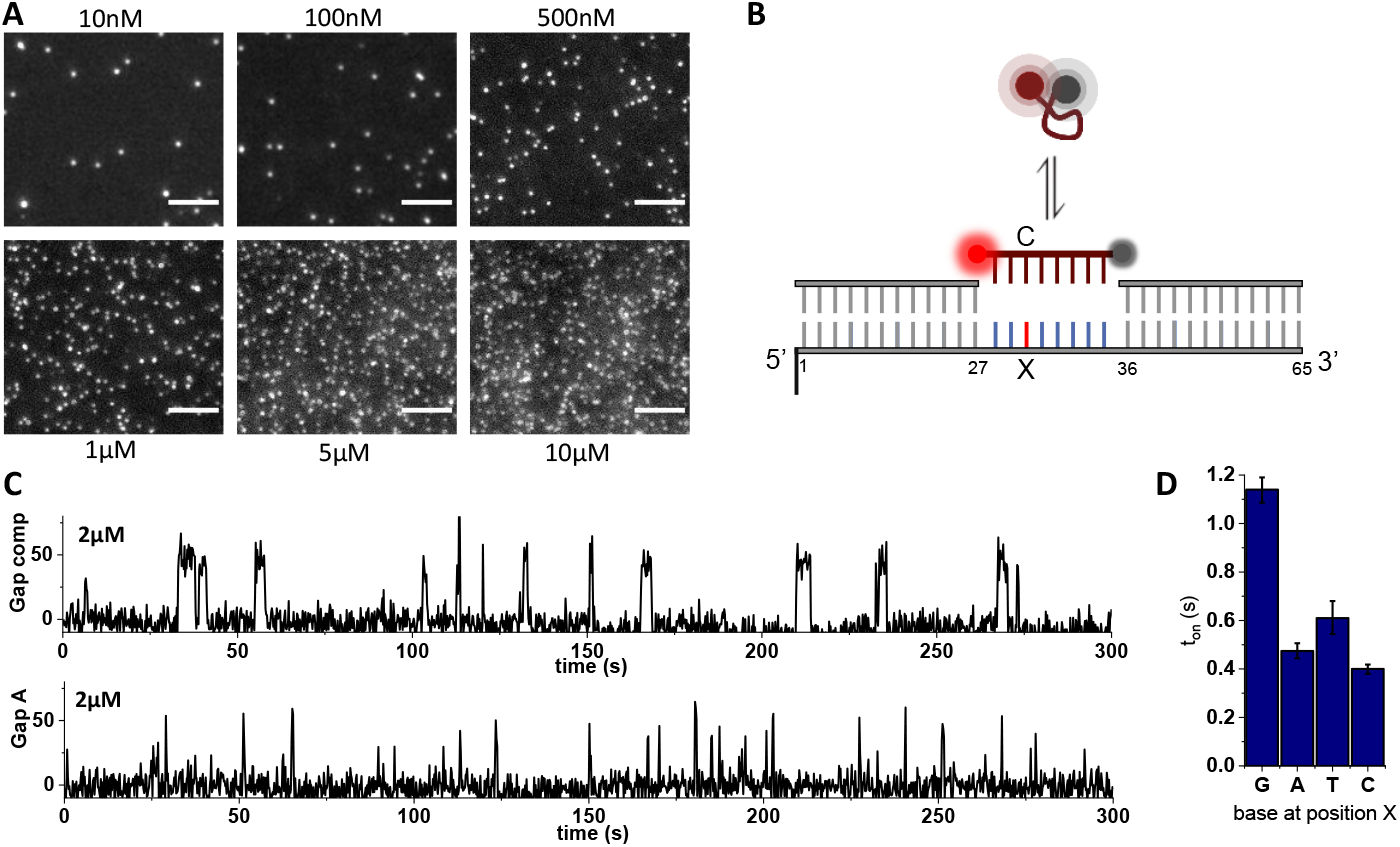
Single-molecule experiments at high concentrations. **A**: Single frame images of the 18 nt probe (described in Figure 4) at various concentrations binding to complementary target strands. Individual molecules can be identified at all concentrations up to 10 µM. **B**. Single-molecule assay for mismatch detection and base calling. The binding kinetics of fluorogenic 8 nt probes will distinguish the fully matched sequence from a mismatch at position labelled with X. **C**. Intensity-vs-time traces of an 8 nt probe binding to a target at 2 µM, top: fully complementary target, bottom A:C mismatch at position X. **D**. The duration of binding events extracted from time traces at 2 µM probes for all for possible bases at position X. Lifetimes were fitted accounting for the camera frame rate and error bars are 95 % confidence intervals determined by bootstrapping.

Operating at high concentrations is particularly important in single-molecule experiments which rely on the accumulation of statistics from individual molecules, and where, without a sufficient on-rate, this process can be long and tedious.

To demonstrate our capability to perform single-molecule experiments at concentrations in the micromolar range, we explored extension of an assay capable of detecting single-base mismatches in a target structure via the hybridisation kinetics of a fluorogenic probe^16^ (Figure 5B). A fully matched target will lead to a more stable interaction with the probe, compared to one with a de-stabilising, single-base mismatch. We performed the assay using 2 µM of fluorogenic probe and achieved a high SNR that allows for robust detection of binding events (Figure 5C). The dwell times of all binding events were extracted for all four possible bases at the position labelled with X, and clearly show the more stable interaction of the probe with the fully complementary target with G at the position X, which leads to longer bound times. The single mismatch reduced the interaction lifetime by a factor of ∼2, allowing a clear distinction of target sequences purely based on the hybridisation kinetics.^16^ Operating at 2 µM has allowed us to accumulate 1000 -3000 fitted binding events within an imaging time of 5 min per field-of-view, averaging between 10 - 20 events per evaluated molecule. Use of a higher concentration of the fluorogenic probe accelerated the process of distinguishing between complementary and mismatch sequences by collecting significant number of binding events in a shorter span of time due to enhanced on-rate at the elevated probe concentration.

### Fast super-resolution imaging with 6-nt imagers in DNA-PAINT

To demonstrate a fast and easy implementation of fluorogenic probes into existing experimental protocols, we performed localisation-based super-resolution imaging using DNA-PAINT with a fluorogenic imager. DNA-PAINT allows for imaging of biological samples through the accumulation of localisations from transiently binding, fluorescently labelled imager strands to complementary docking strands bound to the target structure of interest.^39^

Recently, we have developed a protocol for DNA-PAINT-based imaging of the influenza A viral genome.^41^ In this negative-sense, single-stranded RNA virus, the genome consists of eight viral RNA segments, organised in ribonucleoprotein complexes (vRNPs) in which the RNA wraps around oligomerised virus-encoded nucleoproteins (NPs) and is stabilised in this complex by intermolecular interactions. Super-resolution imaging of the viral segments using DNA-PAINT requires the incubation with segment-specific primary probes, which label the segments with many copies of docking sites for a specific DNA-PAINT imager, as previously established^41^ (Figure 6A). In eight subsequent rounds of imaging, all eight different segments can be imaged within the same sample (and the same individual particles). Performing eight rounds of exchange-PAINT is time-consuming and forms the main bottleneck to accessing the heterogeneity within the viral particles at statistically significant levels.

**Figure 6:**
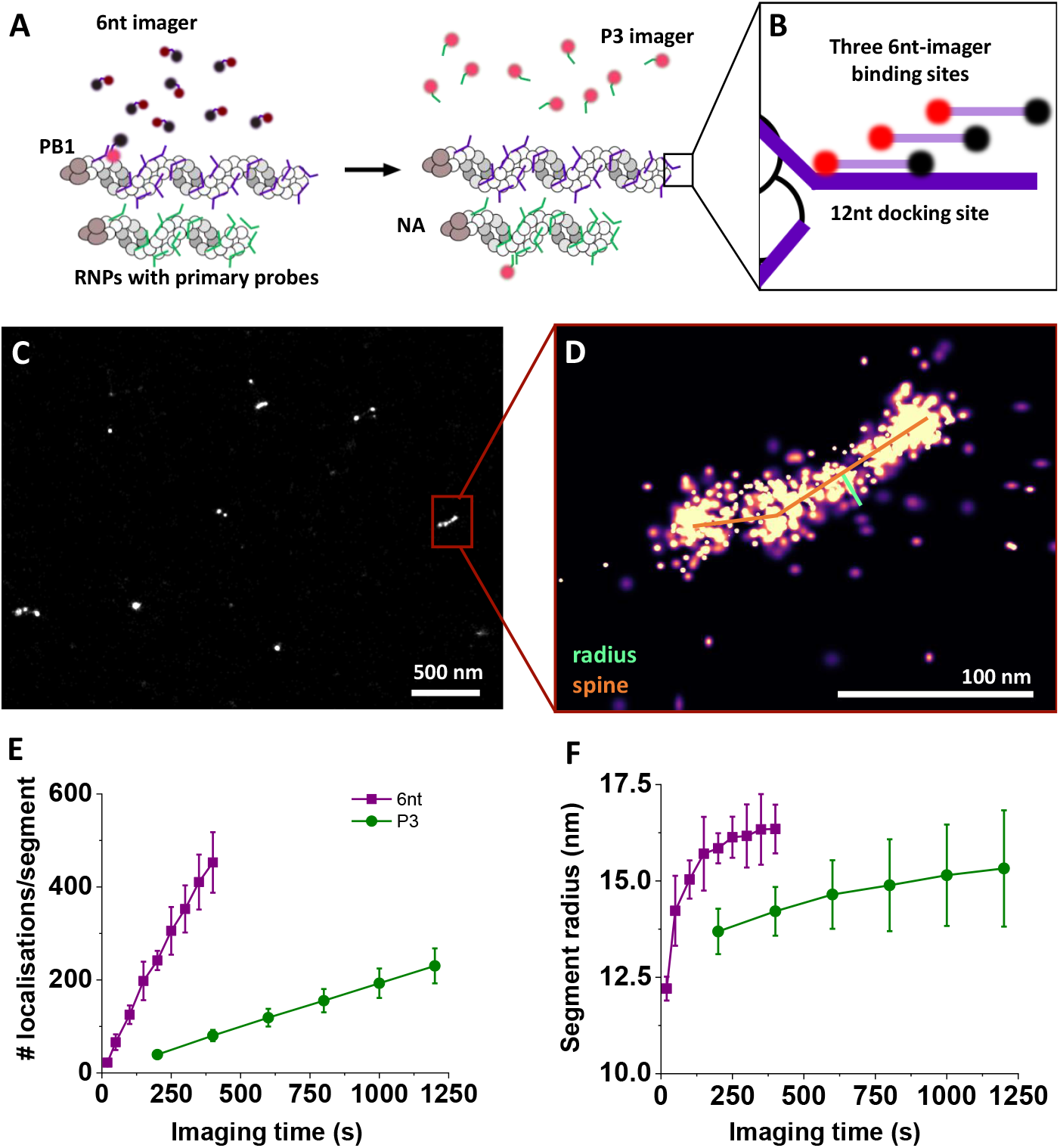
DNA-PAINT on viral particles using 6nt imagers. **A:** General strategy for labelling of the viral genome: RNA-carrying viral segments are hybridised by primary probes which provide the docking sequence for DNA-PAINT imagers. In sequential rounds, the PB1 segment is imaged with the fluorogenic 6 nt imager (purple backbone) followed by the NA segment with the P3 imager. **B:** Arrangement of docking sites for the 6 nt imager. **C:** Reconstructed images of PB1 segment imaged with the 6nt imager. **D:** The zoom-in illustrates the fitted spine and radius. **E:** Localisations per Segment for both imagers. **F:** Segment radius, fitted to the extracted segments. Values are mean and standard deviation of two (6 nt) or three (P3) independent experiments.

Much like other SMLM techniques, the acquisition of a single reconstructed image is time-consuming, mainly because it is the experimental environment which determines imaging speed in DNA-PAINT. There are two key factors to consider: the hybridisation kinetics of imagers to the docking strand and the need for motion blur of unbound imagers, which is required for precise localisation of only target-bound imagers. Both factors limit the imager concentration to the picomolar to low nanomolar range.^39,48,56^ Fluorogenic imagers can help to address both points: through them being essentially dark when unbound, the need for motion blur is significantly reduced, so shorter exposure times can be implemented without detecting unbound imagers. Further, speeding up the actual kinetics of the hybridisation/de-hybridisation cycle requires fast off-rates (i.e. shorter sequences) as well as faster on-rates, facilitated by higher imager concentrations.

Recent efforts to achieve faster or higher-quality imaging include use of fluorogenic imager strands,^34–36^ focusing on the development of specific sequences with mismatches or use of a limited set of dyes to achieve fluorogenicity. However, our fluorogenic approach allows us to create very short imager strands, which can operate at high concentrations and speed up imaging even further.

To demonstrate these capabilities, we employed fluorogenic 6 nt long imagers in imaging viral RNA segments. We imaged two of the eight segments in our experiments: the PB1 segment with probes complementary to the 6-nt imager, and (for comparison) the NA segment with probes complementary to the P3 imager sequence used in the original study (Figure 6A).^41^

The 6nt imager sequence (5’-ATTO643-TGGTGG-BMNQ1-3’) was adopted from published Imager sequences^56^ without further sequence engineering. To maximise the on-rate of imager binding, the primary probes carrying the docking site for the 6-nt imager contained a multiplexed docking sequence (extension: -CCACCACCACCA-), creating three possible binding sites for the imager (a concept first introduced in Strauss et al.^48^, Figure 6B). The fluorogenic properties of the 6-nt imager allowed raising the imager concentration to 20 nM and shorten the exposure time to 20 ms (compared to 5 nM and 200 ms for the P3 imager)^41^ without significant increase in the background levels.

We observe levels of co-localisation of the two segments (imaged with different imagers) which are expected based on the prevalence of each segment type^41^ when imaging the same field of view, and see very few clustered localisations in a negative control sample (a sample without primary probes carrying the docking site for our 6nt imager strand; see Fig S2). These results establish that our signal is specific, and that binding sites are accessible to the doubly-labelled 6nt imager.

We observe more localisations uniformly spaced out across the FOV using the 6-nt imager compared to the P3, which may reflect non-specific surface interactions of the imagers. When imaging at longer exposure times, these very short events are not picked up (or are very dim and can easily be filtered), but when imaging at 20 ms, they are noticeable. To extract the segments, we perform a cluster analysis (see *Methods*) and implemented an additional filtering step in the 6-nt imager data to extract only those clusters with a high core sampling ratio (i.e. high number of central localisations), which originate from true viral segments (Figure 6C).

The observed clusters of localisations can be fitted with a spine along the long axis of the clusters, and a radius (median distance between boundary and spine), both of which report on the physical dimensions of the vRNPs (Figure 6C, D). The fitted spine length of the NA segments imaged with P3 exhibits a wide distribution ranging up to ∼150 nm, potentially caused by projecting the segments oriented in 3D onto the imaging plane.^41^ The segment radius follows a much narrower distribution with a mean of ∼15 – 20 nm, in agreement with previous estimates. ^41,57,58^ We chose the segment radius as the appropriate physical parameter of the viral segments to test our 6-nt imagers and compare to the previous protocol with non-fluorogenic imagers.

Using the 6-nt fluorogenic imagers, we performed DNA-PAINT imaging at 20-ms exposure times (a factor of 10 smaller than the previous protocols). The higher imager concentration (20 nM) increased the on-rate of the 6-nt imager to 1.19 s^-1^ (± 0.02 s^-1^, calculated from localisations per cluster after filtering, Figure 6E), ∼6-fold larger than observed for the P3 imager (0.196 ± 0.001 s^-1^). During image acquisition, the segment sampling increases steadily, and fits of the segment radii converge eventually to their true value (within the error margins affected by localisation precision) of 16 nm (± 1 nm) for the PB1 segment imaged by the 6-nt imager and 13 nm (± 0.5 nm) for the NA segment imaged by P3 (Figure 6F). Notably, the radii could be extracted after imaging of ∼100 binding events per segment, which required ∼800-1000 s of imaging with the P3 imager, and only ∼150 s of imaging with the 6-nt imager (Figure 6E).

In summary, we demonstrated super-resolution imaging of viral genome segments with a fluorogenic 6-nt imager strand. Through increasing the imager concentration whilst shortening the exposure time, the imaging process could be sped up significantly (by a factor of 6). We utilise the key tuning screws of the fluorogenicity to facilitate sufficient de-quenching in the hybridised state of the 6-nt imager, overcoming limitations of previous fluorogenic strategies failing at short probe lengths.

## Conclusion

We have conducted an in-depth characterisation of the fluorogenicity in self-quenching ssDNA probes (terminally labelled with F and Q), evaluating the effect of probe length, R_0_ of the F/Q pair, and transient secondary interactions on fluorogenicity.

Previously, optimising for the highest degree of quenching in the unbound state led to the conclusion that a minimum of ∼15-nt distance had to be incorporated between dye and quencher to allow for sufficient de-quenching for single-molecule detection. ^34,36^ However, by focusing on the principles guiding the de-quenching efficiency, we constructed highly fluorogenic probes of various lengths, down to 5 nt. We also devised strategies which can increase quenching (and thereby fluorogenicity) in longer probes.

Our findings can easily be translated to other **spectral regions**. Although we chose the red dye ATTO647N for our characterisation, we have also performed experiments with ATTO643. The only spectral or dye-specific property in our design is the R_0_ of the F/Q pair, which can easily be determined for most F-Q combination by evaluating available spectral data, independent of the spectral region of interest. Previous approaches on short (8-12 nt) ssDNA fluorogenic probes have used a self-quenching approach, whereby two terminally fluorophores on the probe would contact-quench in solution.^16,35,37^ This approach is, however, limited to very specific fluorophores (e.g. ATTO647N and ATTO655), and does not easily translate to other spectral regions.

Possible contributions from contact-quenching in the unbound state when using F-Q pairs (not addressed in our study) will further improve the FF, as the unbound state will be further darkened without affecting the de-quenched fluorescence significantly (given the same R_0_). Such contributions could be optimised by finding the most efficient Qs which exhibit a similar R_0_ with the F of choice. Because R_0_ values are measurable and -to a large degree - predictable, Qs can be pre-selected for their R_0_ and then only a subset of suitable candidates tested for the best contact quenchers – a property much more difficult to predict without experimental testing.

The **probe length** in our design is fully flexible within the regions interesting for hybridisation kinetics spanning the timescales of most experiments (5 - 25 nt), and via the introduction of secondary structures, even longer designs are possible. We are, however, not relying on specific structural motifs, giving the experimentalist complete freedom over the sequences used.

Overall, this allows us to generate plug-and-play level solutions to introduce fluorogenicity into experimental designs.

To demonstrate the ease of implementation of such solutions into existing protocols, we performed **DNA-PAINT** imaging on influenza A viral segments with a **6-nt long, fluorogenic imager** and showed that we can recover the physical dimensions accurately, and 6-fold faster than in the initial viral imaging protocol using a non-fluorogenic imager. DNA-PAINT uses point-spread-function fitting to localise every emitter, a process much more sensitive to background than, e.g., the SMF assays we performed in this study. Thus, fluorogenic imagers were required to perform good-quality DNA-PAINT imaging even when operating below the technical TIRF concentration barrier of ≈ 100 nM. More generally, the background reduction will improve the SNR at any concentration, making it useful in single-molecule measurements at all concentration ranges.

The introduction of fluorogenicity into virtually any ssDNA label will allow a wide range of studies previously inaccessible due to lack of signal specificity, or high levels of background fluorescence. Concentrations in the micromolar range (as we demonstrated in our SMF assay) allow to speed up reaction and binding kinetics of the ssDNA probes.

Indirectly, fluorogenicity opens experimental avenues to circumvent and address old problems and questions of single-molecule fluorescence experiments. One such application allowed us to implement virtually bleaching-free single-molecule FRET-measurements through the constant exchange of fluorophores delivered on fluorogenic ssDNA r-labels at high concentrations.^37^

Through the high signal specificity of fluorogenic probes (unbound probes can be virtually dark), our fluorogenic probes will be of great advantage in live-cell single-molecule imaging and tracking applications, extending the current state-of-the-art where ssDNAs for fluorophore delivery are predominantly used in fixed cells that require washing of unbound probes. Such a capability will pave the way for direct imaging of biomolecular structure and interactions in vivo, opening a plethora of exciting biological applications and assays.

## Authors Contributions

Mirjam Kümmerlin: Conceptualisation, Project administration, Methodology, Investigation, Formal analysis, Visualisation, Writing original draft; Qing Zhao: Formal Analysis, Software, Visualisation; Jagadish Hazra: Methodology; Christof Hepp: Methodology; Alison Farrar: Methodology, Investigation, Formal analysis, Visualisation; Piers Turner: Software; Achillefs N. Kapanidis: Conceptualisation, Project administration, Writing – reviewing and editing, Funding Acquisition

## References

1. Miller, H., Zhou, Z., Shepherd, J., Wollman, A. J. M. & Leake, M. C. Single-molecule techniques in biophysics: a review of the progress in methods and applications. Reports on Progress in Physics 81, 24601 (2017).

2. Xie, X. S., Choi, P. J., Li, G. W., Nam, K. L. & Lia, G. Single-Molecule Approach to Molecular Biology in Living Bacterial Cells. Annu Rev Biophys 37, 417–444 (2008).

3. Lelek, M. et al. Single-molecule localization microscopy. Nature Reviews Methods Primers 1, 39 (2021).

4. Löschberger, A. et al. Super-resolution imaging visualizes the eightfold symmetry of gp210 proteins around the nuclear pore complex and resolves the central channel with nanometer resolution. J Cell Sci 125, 570–575 (2012).

5. Hess, S. T., Girirajan, T. P. K. & Mason, M. D. Ultra-High Resolution Imaging by Fluorescence Photoactivation Localization Microscopy. Biophys J 91, 4258–4272 (2006).

6. Betzig, E. et al. Imaging Intracellular Fluorescent Proteins at Nanometer Resolution. Science (1979) 313, 1642–1645 (2006).

7. Pertsinidis, A., Zhang, Y. & Chu, S. Subnanometre single-molecule localization, registration and distance measurements. Nature 466, 647–651 (2010).

8. Rust, M. J., Bates, M. & Zhuang, X. Sub-diffraction-limit imaging by stochastic optical reconstruction microscopy (STORM). Nat Methods 3, 793–796 (2006).

9. Gelles, J., Schnapp, B. J. & Sheetz, M. P. Tracking kinesin-driven movements with nanometre-scale precision. Nature 331, 450–453 (1988).

10. Schmidt, T., Schutz, G. J., Baumgartner, W., Gruber, H. J. & Schindler, H. Imaging of single molecule diffusion. Proceedings of the National Academy of Sciences 93, 2926–2929 (1996).

11. Tinnefeld, P. Breaking the concentration barrier. Nat Nanotechnol 8, 480–482 (2013).

12. Holzmeister, P., Acuna, G. P., Grohmann, D. & Tinnefeld, P. Breaking the concentration limit of optical single-molecule detection. Chem. Soc. Rev. 43, 1014–1028 (2014).

13. White, D. S., Smith, M. A., Chanda, B. & Goldsmith, R. H. Strategies for Overcoming the Single-Molecule Concentration Barrier. ACS Measurement Science Au 3, 239–257 (2023).

14. Magaki, S., Hojat, S. A., Wei, B., So, A. & Yong, W. H. An Introduction to the Performance of Immunohistochemistry. in 289–298 (2019). doi:10.1007/978-1-4939-8935-5_25.

15. Cook, A., Walterspiel, F. & Deo, C. HaloTag-Based Reporters for Fluorescence Imaging and Biosensing. ChemBioChem 24, (2023).

16. Andrews, R. et al. Transient DNA binding to gapped DNA substrates links DNA sequence to the single-molecule kinetics of protein-DNA interactions. bioRxiv 2022.02.27.482175 (2022) doi:10.1101/2022.02.27.482175.

17. Peng, S., Wang, W. & Chen, C. Breaking the Concentration Barrier for Single-Molecule Fluorescence Measurements. Chemistry – A European Journal 24, 1002–1009 (2018).

18. van Oijen, A. M. Single-molecule approaches to characterizing kinetics of biomolecular interactions. Curr Opin Biotechnol 22, 75–80 (2011).

19. Puchkova, A. et al. DNA Origami Nanoantennas with over 5000-fold Fluorescence Enhancement and Single-Molecule Detection at 25 μM. Nano Lett 15, 8354–8359 (2015).

20. Lukinavičius, G. et al. Fluorogenic probes for live-cell imaging of the cytoskeleton. Nat Methods 11, 731–733 (2014).

21. Kozma, E. & Kele, P. Fluorogenic probes for super-resolution microscopy. Org Biomol Chem 17, 215–233 (2019).

22. Sharonov, A. & Hochstrasser, R. M. Wide-field subdiffraction imaging by accumulated binding of diffusing probes. Proceedings of the National Academy of Sciences 103, 18911–18916 (2006).

23. Flors, C. DNA and chromatin imaging with super-resolution fluorescence microscopy based on single-molecule localization. Biopolymers 95, 290–297 (2011).

24. Livak, K. J., Flood, S. J. A., Marmaro, J., Giusti, W. & Deetz, K. Oligonucleotides with fluorescent dyes at opposite ends provide a quenched probe system useful for detecting PCR product and nucleic acid hybridization. PCR Methods Appl 4, 357–362 (1995).

25. Lee, L. G., Connell, C. R. & Bloch, W. Allelic discrimination by nick-translation PCR with fluorogenic probes. Nucleic Acids Res 21, 3761–3766 (1993).

26. Holland, P. M., Abramson, R. D., Watson, R. & Gelfand, D. H. Detection of specific polymerase chain reaction product by utilizing the 5’ 3’ exonuclease activity of Thermus aquaticus DNA polymerase. Proceedings of the National Academy of Sciences 88, 7276–7280 (1991).

27. Tyagi, S. & Kramer, F. R. Molecular Beacons: Probes that Fluoresce upon Hybridization. Nature Biotechnology 1996 14:3 14, 303–308 (1996).

28. Tyagi, S., Bratu, D. P. & Kramer, F. R. Multicolor molecular beacons for allele discrimination. Nat Biotechnol 16, (1998).

29. Johansson, M. K., Fidder, H., Dick, D. & Cook, R. M. Intramolecular Dimers: A New Strategy to Fluorescence Quenching in Dual-Labeled Oligonucleotide Probes. J Am Chem Soc 124, 6950–6956 (2002).

30. Dubertret, B., Calame, M. & Libchaber, A. J. Single-mismatch detection using gold-quenched fluorescent oligonucleotides. Nature Biotechnology 2001 19:4 19, 365–370 (2001).

31. Nazarenko, I. Multiplex quantitative PCR using self-quenched primers labeled with a single fluorophore. Nucleic Acids Res 30, 37e–337 (2002).

32. Marras, S. A. E., Kramer, F. R. & Tyagi, S. Efficiencies of fluorescence resonance energy transfer and contact-mediated quenching in oligonucleotide probes. Nucleic Acids Res 30, (2002).

33. Le Reste, L., Hohlbein, J., Gryte, K. & Kapanidis, A. N. Characterization of dark quencher chromophores as nonfluorescent acceptors for single-molecule FRET. Biophys J 102, 2658–2668 (2012).

34. Chung, K. K. H. et al. Fluorogenic DNA-PAINT for faster, low-background super-resolution imaging. Nat Methods 19, 554–559 (2022).

35. Kessler, L. F. et al. Self-quenched Fluorophore Dimers for DNA-PAINT and STED Microscopy. Angew Chem Int Ed Engl 62, (2023).

36. Schueder, F. & Bewersdorf, J. Highly Multiplexed Imaging with Speed and Fluorogenic DNA-PAINT. Microscopy and Microanalysis 29, 1069–1069 (2023).

37. Kümmerlin, M., Mazumder, A. & Kapanidis, A. N. Bleaching-resistant, Near-continuous Single-molecule Fluorescence and FRET Based on Fluorogenic and Transient DNA Binding. ChemPhysChem 24, e202300175 (2023).

38. Vietz, C., Lalkens, B., Acuna, G. P. & Tinnefeld, P. Synergistic Combination of Unquenching and Plasmonic Fluorescence Enhancement in Fluorogenic Nucleic Acid Hybridization Probes. Nano Lett 17, 6496–6500 (2017).

39. Schnitzbauer, J., Strauss, M. T., Schlichthaerle, T., Schueder, F. & Jungmann, R. Super-resolution microscopy with DNA-PAINT. Nat Protoc 12, 1198 (2017).

40. Woody, M. S., Lewis, J. H., Greenberg, M. J., Goldman, Y. E. & Ostap, E. M. MEMLET: An Easy-to-Use Tool for Data Fitting and Model Comparison Using Maximum-Likelihood Estimation. Biophys J 111, 273–282 (2016).

41. Hepp, C., Zhao, Q., Robb, N., Fodor, E. & Kapanidis, A. Structural Analysis of the Influenza Genome by High-Throughput Single-Virion DNA-PAINT. To be submitted to biorxiv. (Jan 2025).

42. Etheridge, T. J., Carr, A. M. & Herbert, A. D. GDSC SMLM: Single-molecule localisation microscopy software for ImageJ. Wellcome Open Res 7, 241 (2022).

43. Ma, H., Chen, M., Nguyen, P. & Liu, Y. Toward drift-free high-throughput nanoscopy through adaptive intersection maximization. Sci Adv 10, (2024).

44. Wang, Y., Gu, Y. & Shun, J. Theoretically-Efficient and Practical Parallel DBSCAN. in Proceedings of the 2020 ACM SIGMOD International Conference on Management of Data 2555–2571 (ACM, New York, NY, USA, 2020). doi:10.1145/3318464.3380582.

45. Ridler, T. W. & Calvard, S. Picture thresholding using an iterative selection method. IEEE Trans Syst Man Cybern SMC-8, 630–632 (1978).

46. Verwer, B. J. H. Improved metrics in image processing applied to the Hilditch skeleton. in [1988 Proceedings] 9th International Conference on Pattern Recognition 137–142 (IEEE Comput. Soc. Press). doi:10.1109/ICPR.1988.28189.

47. Jungmann, R. et al. Multiplexed 3D cellular super-resolution imaging with DNA-PAINT and Exchange-PAINT. Nat Methods 11, 313–318 (2014).

48. Strauss, S. & Jungmann, R. Up to 100-fold speed-up and multiplexing in optimized DNA-PAINT. Nat Methods 17, 789–791 (2020).

49. Linko, V. et al. One-step large-scale deposition of salt-free DNA origami nanostructures. Sci Rep 5, 15634 (2015).

50. Kielar, C. et al. On the Stability of DNA Origami Nanostructures in Low-Magnesium Buffers. Angewandte Chemie International Edition 57, 9470–9474 (2018).

51. Chen, H. et al. Ionic strength-dependent persistence lengths of single-stranded RNA and DNA. Proc Natl Acad Sci U S A 109, 799–804 (2012).

52. Zhang, Z., Revyakin, A., Grimm, J. B., Lavis, L. D. & Tjian, R. Single-molecule tracking of the transcription cycle by sub-second RNA detection. Elife 3, (2014).

53. Tyagi, S. & Kramer, F. R. Molecular beacons: probes that fluoresce upon hybridization. Nat Biotechnol 14, 303–308 (1996).

54. Asanuma, H., Fujii, T., Kato, T. & Kashida, H. Coherent interactions of dyes assembled on DNA. Journal of Photochemistry and Photobiology C: Photochemistry Reviews 13, 124–135 (2012).

55. Zadeh, J. N. et al. NUPACK: Analysis and design of nucleic acid systems. J Comput Chem 32, 170–173 (2011).

56. Schueder, F. et al. An order of magnitude faster DNA-PAINT imaging by optimized sequence design and buffer conditions. Nat Methods 16, 1101–1104 (2019).

57. Moeller, A., Kirchdoerfer, R. N., Potter, C. S., Carragher, B. & Wilson, I. A. Organization of the influenza virus replication machinery. Science (1979) 338, 1631–1634 (2012).

58. Arranz, R. et al. The structure of native influenza virion ribonucleoproteins. Science (1979) 338, 1634–1637 (2012).

