## Supplementary Material for "Tunable fluorogenic DNA probes drive fast and high-resolution single-molecule fluorescence imaging"

#### Spectral data for determining the Förster Radius:

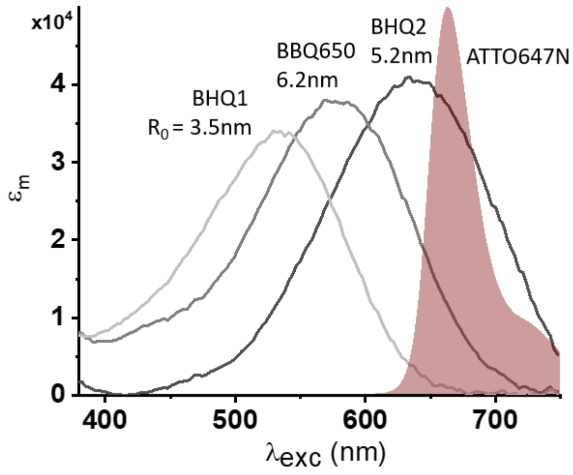


*Figure S1: Determination of the Förster Radius. Quencher absorption spectra and the ATTO647N emission spectrum with calculated Förster Radii. We measured absorption spectra of all Quenchers on the Nanodrop, and used the Fluorophore emission spectra and fluorescence lifetimes provided by the manufacturer.*[1,2] *The Förster radii were then calculated using the FRET-Calc tool.*[3]

#### DNA-PAINT 6nt imager Specificity

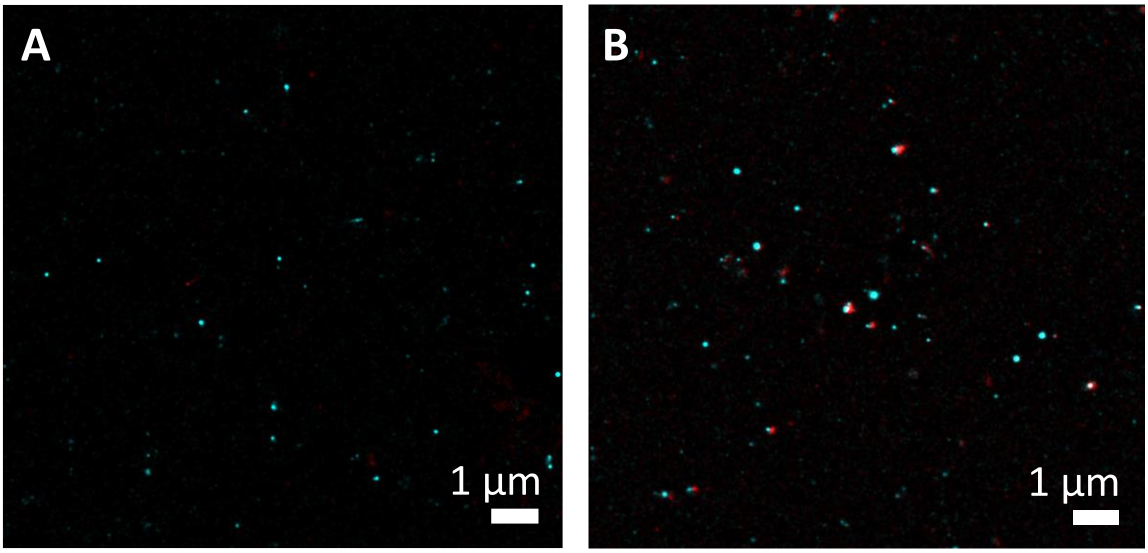


*Figure S2: Control experiments for 6nt DNA-PAINT imaging.* ***A****: Negative control without primary probes (red) shows few localisations compared to the sample with primary probes (cyan).* ***B:*** *P1 imager (Cy3B, Cyan) and 6nt imager (A647N, red) colocalise at viral segments.*

#### Sequences of DNAs

*Table S1 DNA Sequences used in this publication.*

| *Name* | *Sequence 5'-> 3'* | *5' modification* | *3' modification* |
|---|---|---|---|
| ***Characterisation*** | | | |
| *SP_5_A647N_BHQ1* | TTT TT | A647N | BHQ1 |
| *SP_6_A647N_BHQ1* | TTT TTT | A647N | BHQ1 |
| *SP_8_A647N_BHQ1* | TTT GGT TT | A647N | BHQ1 |
| *SP_10_A647N_BHQ1* | TTT GTG GTT T | A647N | BHQ1 |
| *SP_12_A647N_BHQ1* | TTT GTT GGT TTT | A647N | BHQ1 |
| *SP_15_A647N_BHQ1* | TTT GTT GGT TGG TTT | A647N | BHQ1 |
| *SP_20_A647N_BHQ1* | TTT GTT GGT TGG GTT GTT TT | A647N | BHQ1 |
| *SP_25_A647N_BHQ1* | TTT GTT GGT TGG GTT GTG TTG GTT T | A647N | BHQ1 |
| *SP_6_comp_bio* | AAA AAA | biotin | |
| *SP_8_comp_bio* | AAA CCA AA | biotin | |
| *SP_10_comp_bio* | AAA CCA CAA A | biotin | |
| *SP_12_comp_bio* | AAA ACC AAC AAA | biotin | |
| *SP_15_comp_bio* | AAA CCA ACC AAC AAA | biotin | |
| *SP_20_comp_bio* | AAA ACA ACC CAA CCA ACA AA | biotin | |
| *SP_25_comp_bio* | AAA CCA ACA CAA CCC AAC CAA CAA A | biotin | |
| *SP _15_BHQ1* | TTT GTT GGT TGG TTT | A647N | BHQ1 |
| *SP _15_BHQ2* | TTT GTT GGT TGG TTT | A647N | BHQ2 |
| *SP _15_BBQ650* | TTT GTT GGT TGG TTT | A647N | BBQ650 |
| *SP _15_A647N* | TTT GTT GGT TGG TTT | A647N | |
| *SP _18_2ndry* | GCTGCCTCCCGTAGGAGT | A647N | BHQ1 |
| *SP_18_comp_2ndry_bio* | ACTCCTACGGGAGGCAGC | biotin | |
| ***DNA-PAINT*** | | | |
| *P3* | GTAATGAAGA | Cy3B | |
| *6ntl_A643_BMNQ1* | TGGTGG | Atto 643 | BMNQ1 |
| *6ntl_A647N_BHQ1* | TGGTGG | A647N | BHQ1 |
| *comp_R2-6ntl_bio* | TTTCCACCA | biotin | |
| *Primary Probes 6nt_PB1* | CCACCACCACCA-R | | R = comp. PB1 Sequences [4] |
| *Primary Probes P3_NA* | TCTTCATTAC-R | | R= comp. NA Sequences [4] |
| ***Single-molecule assay*** | | | |
| *gap_left* | CCTCATTCTTCGTCCCA**T**TACCATACA | Cy3B | |
| *gap_right* | CGATAATCTGCTGCCTCAGGCTCTTGAC TG | | |
| *seal* | TCCACCGT | A643 | BHQ1 |
| *bottom_A* | CAGTCAAGAGCCTGAGGCAGCAGATTAT CGACGGT**A**GATGTATGGTAATGGGACGA AGAATGAGG | | Biotin |
| *bottom_T* | CAGTCAAGAGCCTGAGGCAGCAGATTAT CGACGGT**T**GATGTATGGTAATGGGACGA AGAATGAGG | | Biotin |
| *bottom_G* | CAGTCAAGAGCCTGAGGCAGCAGATTAT CGACGGTGGATGTATGGTAATGGGACGA AGAATGAGG | | Biotin |
| *bottom_C* | CAGTCAAGAGCCTGAGGCAGCAGATTAT CGACGGT**C**GATGTATGGTAATGGGACGA AGAATGAGG | | Biotin |

### References

[1] ATTO-TEC GmbH, “ATTO643 product information,” **2022**.

[2] ATTO-TEC GmbH, “ATTO647N product Information,” **2022**.

[3] L. Benatto, O. Mesquita, J. L. B. Rosa, L. S. Roman, M. Koehler, R. B. Capaz, G. Candiotto, *Comput Phys Commun* **2023**, *287*, 108715.

[4] C. Hepp, Q. Zhao, N. Robb, E. Fodor, A. Kapanidis, to be submitted to biorxiv, **Jan 2025**.
